# NextflowWorkbench: Reproducible and Reusable Workflows for Beginners and Experts

**DOI:** 10.1101/041236

**Authors:** Jason P. Kurs, Manuele Simi, Fabien Campagne

**Affiliations:** The HRH Prince Alwaleed Bin Talal Bin Abdulaziz Alsaud Institute for Computational Biomedicine, Weill Cornell Medicine, New York, NY, United States of America; Clinical Translational Science Center, Weill Cornell Medicine, New York, NY, United States of America; Department of Physiology and Biophysics, Weill Cornell Medicine, New York, NY United States of America

**Keywords:** Workflows, Pipelines, Reproducibility, Docker, Language Workbench, JetBrains MPS

## Abstract

Computational workflows and pipelines are often created to automate series of processing steps. For instance, workflows enable one to standardize analysis for large projects or core facilities, but are also useful for individual biologists who need to perform repetitive data processing. Some workflow systems, designed for beginners, offer a graphical user interface and have been very popular with biologists. In practice, these tools are infrequently used by more experienced bioinformaticians, who may require more flexibility or performance than afforded by the user interfaces, and seem to prefer developing workflows with scripting or command line tools. Here, we present a workflow system, the NextflowWorkbench (NW), which was designed for both beginners and experts, and blends the distinction between user interface and scripting language. This system extends and reuses the popular Nextflow workflow description language and shares its advantages. In contrast to Nextflow, NextflowWorkbench offers an integrated development environment that helps complete beginners get started with workflow development. Auto-completion helps beginners who do not know the syntax of the Nextflow language. Reusable processes provide modular workflows. Programmers will benefit from unique interactive features that help users work more productively with docker containers. We illustrate this tool with a workflow to estimate RNA-Seq counts using Kallisto. We found that beginners can be taught how to assemble this workflow in a two hours training session. NW workflows are portable and can execute on laptop/desktop computers with docker, on a lab cluster, or in the cloud to facilitate training. NextflowWorkbench is open-source and available at http://workflow.campagnelab.org.

## INTRODUCTION

A computational workflow or pipeline is a description of a series of computational steps connected to each other. Each step accepts one or more input(s) and transforms input(s) to produce one or more output(s). Computational workflows are used in many engineering and scientific domains, but are particularly useful in fields such as bioinformatics where analysis activities are repetitive and benefit from being automated. Workflows can be represented as diagrams and their steps followed manually, but many solutions have been developed to represent workflows electronically and automate their execution.

Automated workflow systems include a way for a user to edit the formal representation of the workflow, and its component steps, as well as a runtime system to execute specific workflows. Two broad families of workflow systems have been developed and are still in use today.

The first category is workflows with graphical user interfaces, which often represent workflows as connected components on a 2D diagram. There is a long history for such tools, but Galaxy and GenePattern are well known examples in Bioinformatics Giardine et al. [2005], Reich et al. [2006]. Workflow systems with graphical user interfaces are favored by beginners, or by educators who need to teach beginners (see this thread, for instance, where these arguments have been made by others https://www.biostars.org/p/50034/).

The second category of workflow systems is based on programming and scripting languages. A workflow is expressed in a declarative or imperative language, or a combination of both. An example of declarative language is the makefile input of the make/gmake tool available in UNIX, which was initially developed to automate the compilation of programs, but has been used as well to implement workflows. An example of imperative workflow is when bioinformaticians develop a collection of scripts and execute these scripts to implement different pipelines.

While useful, many of these systems fail to fulfill needs that are common in bioinformatics. For instance a makefile cannot easily be run in a distributed manner on multiple nodes of a cluster to parallelize the processing of large collections of data. Scripts can be run in parallel on a cluster with tools like GNU parallel or grid schedulers (e.g., Sun Grid Engine or SLURM), but a recurrent problem with scripts is that most of them are not easily portable to new environments. Indeed, most scripts are written to assume a specific location for the programs that they use (either the script expects a program to be in the PATH, or the script contains a dependency on some location defined in the system where the script was developed). When a dependency is not available in the location assumed by the script, the script fails and the user needs to resolve the issue before retrying execution. The problem of managing dependencies for scripts is often described as “dependency hell”, a term which many people believe adequately describes the practical difficulties of getting scripts to run on other systems than where they have been developed.

Several groups have recently recognized these problems and have developed improved solutions targeted at bioinformaticians. Recent developments include BigDataScript Cingolani et al. [2015] and Nextflow Di Tommaso et al. [2014] (http://nextflow.io). These solutions address scalability and portability problems and are useful to users with programming and/or scripting experience.

We recently asked the question of whether we could design a hybrid between scripting and user interface workflow systems. Such a hybrid system would make it possible for beginners and experts to collaborate more closely by using the same platform to represent and execute workflows. This manuscript describes the NextflowWorkbench, a workflow platform that addresses this question. We developed NextflowWorkbench with Language Workbench Technology implemented in the JetBrains MPS system (jetbrains.com/mps) Dmitriev [2004], Campagne [2014, 2015], Simi and Campagne [2014], Benson and Campagne [2015].

This platform leverages Nextflow (see Di Tommaso et al. [2014] and http://nextflow.io), a workflow language developed for users familiar with the command line and scripting. NW shares most of the advantages of Nextflow, and adds features required for modularity and reuse. Importantly, NW also offers interactive features designed to guide new users who are not familiar with the syntax of the Nextflow language. We report on the design of NextflowWorkbench and on our experience teaching this new platform to biologists and clinicians with no prior scripting experience.

## METHODS

We have used the MPS Language Workbench (http://jetbrains.com/mps), as also described in Campagne [2014] and Campagne [2015]. For an introduction to Language Workbench Technology (LWT) in the context of bioinformatics see Simi and Campagne [2014] and Benson and Campagne [2015], Campagne and Simi [2015] in the context of data analysis. Methods for this study are similar to those described in Campagne et al. [2015].

### Language Design

JPK designed and developed the core MPS languages of NextflowWorkbench during a three month summer internship in the Campagne laboratory. These core languages include functionality to represent Processes, Workflow and execution Scripts. Additional developments were conducted by MS and FC to extend these languages and add docker (https://docker.com) and GobyWeb functionality. Full language development logs are available on the GitHub code repository (https://github.com/CampagneLaboratory/NextflowWorkbench). Briefly, we designed abstractions to represent Nextflow scripts in a modular fashion (decoupling Processes from Workflows). These abstractions were implemented with the structure, editor, constraints and typesystem aspects of MPS languages (described in Campagne [2014]). Importantly, the typesystem aspect makes it possible to typecheck a workflow as it is being developed and provide feedback to the developer. Nextflow scripts are generated from nodes of the languages using the MPS textgen aspect (Campagne [2014]).

### Workflow Execution

Workflows can be executed directly from within the MPS LW. Execution is supported on the developers’ machine (for Linux and Mac OS platforms only, since Nextflow does not run on Windows), on a remote Linux node, where one of the execution mechanisms supported by Nextflow must be available (e.g., Sun Grid Engine, SLURM, Apache Ignite), or in the Cloud (Google Cloud platform is supported at this time, see cluster provisioning). This capability was implemented with Run Configurations (see Campagne [2015], Chapter 5). Executing a workflow on a lab cluster or in the cloud is done the same way after configuring the IP address or hostname of the cluster front-end, and the necessary credentials to SSH onto this server.

### Cluster Provisioning in the Cloud

We have developed automated methods to provision a cluster in the cloud. Briefly, we use the Google Genomics branch of elasticluster (http://googlegenomics.readthedocs.org/en/latest/use_cases/setup_gridengine_cluster_on_compute_engine/) to configure a cluster with one front-end and *n* nodes, where *n* is configurable by the end user of NW. Because Elasticluster has a complicated installation, which most beginners may not be able to complete, we prepared a docker image which contains all the necessary tools and customization needed to create a cluster. NW offers a language with a user interface to configure the parameters of a cluster (e.g., number of compute nodes, type of node, how much storage each node should have, etc.). Pressing a button on this user interface (see Figure 8) executes the appropriate script inside the elasticluster container and creates or destroys a cluster.

### Scripts

To implement scripts as text with auto-completion, we used the MPS RichText plugin (developed and distributed by members of the MBEDDR project Voelter et al. [2012]). The plugin implements the approach described in Voelter [2013].

### Documentation

Project documentation has been developed with LTEX and the Editor2PDF language and plugin (https://github.com/CampagneLaboratory/Editor2PDF). Complete documentation is available at Kurs and Campagne [2015].

## RESULTS

### A Workflow System for Beginners and Experts

In this study, we designed a workflow system aimed at the full spectrum of workflow users, from beginners to computational experts. Figure 1 presents the advantages of this new workflow system across its intended spectrum of users. The following results section describes the design of this system and the innovations introduced to help with the development and maintenance of reproducible and high-performance workflows. This section also addresses the question of whether this system can be taught effectively to beginners with no programming or command line experience.

**Figure 1.**
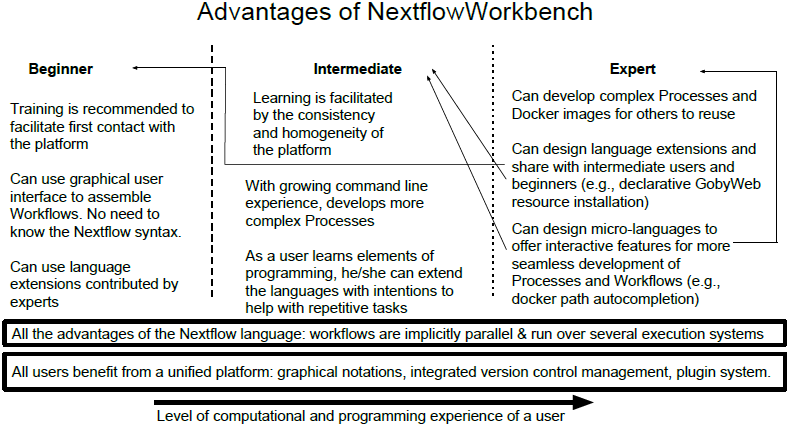
Advantages of the NextflowWorkbench workflow system across its intended user spectrum.

### Evaluation of Workflow Systems

We selected Nextflow as the target language for the NextflowWorkbench platform after comparing three systems with similar goals. The comparison included BigDataScript (http://pcingola.github.io/BigDataScript/, Cingolani et al. [2015]), Nextflow (http://nextflow.io, Di Tommaso et al. [2014]) and the Swift language (http://swift-lang.org/, Wilde et al. [2011]). These systems were selected for evaluation because they support parallel execution of workflows on multi-node clusters and provide a reasonable level of abstraction to express workflows.

To compare these systems, two evaluators tried to implement a simple analysis pipeline with each of them. One evaluator was an experienced software engineer with decades of programming and scripting experience (MS). The second evaluator (JPK) was a sophomore undergraduate student with intermediate level programming and scripting experience. Both evaluators completed the implementation of the test pipeline with BigDataScript and Nextflow. The more experienced evaluator developed a partial prototype workflow with Swift, but reported that locating information in the online documentation was tedious and that the semantic of the language was far from intuitive.

This short evaluation revealed a number of characteristics, advantages and drawbacks of the three systems:

**Swift** requires that all programs used in the workflow are already installed on each machine of a cluster. A text file binds the name of a program used in the Swift script to the path of this program on the machine. This requirement means that Swift assumes that dependencies of a workflow are pre-installed and fixed, and makes no effort to facilitate the installation of programs and dependencies on cluster nodes.

**BigDataScript** was presented in detail in Cingolani et al. [2015]. During our evaluation, we noticed a bug report in the forum that indicated a major error in dataflow analysis of the script by the BigDataScript compiler/interpreter (see https://groups.google.eom/forum/#Itopic/bigdatascript-users/r7rQ03LBYIc. The magnitude of this error suggested to us that BigDataScript was not, at the time, a robust language for developing workflows.

**Nextflow** performed as expected and the evaluators found the documentation sometimes difficult to follow, but overall fairly complete and sufficient. The language offers direct support for running steps of a workflow— called Processes in the Nextflow language— inside a docker container. This feature is extremely useful to develop workflows that can execute on other systems without tedious dependency installation Di Tommaso et al. [2015]. Negatives were the lack of language modularity, making it impossible to develop libraries of reusable processes and the difficulty of knowing at first glance ‐at least for beginners who do not know the language well— what type of data is exchanged between processes.

The evaluators concluded that of the three systems, Nextflow was the more promising system for representing workflows as scripts.

### Requirements for an Improved Workflow Language

Following up this evaluation, we decided to design a variant of the Nextflow language that would directly address the limitations that our evaluation had identified. Specifically, we wanted:

- A modular workflow language that would make it straightforward to reuse processes developed by others in new workflows. Modularity can enable experts to develop processes and share these processes with beginners.
- An explicitly typed workflow language. We believe that an explicitly typed language makes it more obvious to beginners what data are expected as input to a process and what data will be produced as output. Coupled with a mechanism to check type compatibility (a type system) at runtime and highlight type errors, explicit types make it easier for beginners to develop correct workflows. A type system is also useful to experts because it highlights errors that could be missed and only become apparent when trying to execute the workflow.
- A language that does not require the user to know or remember its precise syntax to use it. Such languages can be built with language workbench technology to provide auto-completion that guides beginners and experts. We have shown the effectiveness of these approaches to help beginners in the MetaR project (Campagne et al. [2015]) and were curious to find out if they could also help with workflow development.

### Process

A NexflowWorkbench Process is illustrated in Figure 2. A process consists of a set of inputs, a set of outputs, and a script, which implements the processing of inputs to generate outputs.

**Figure 2.**
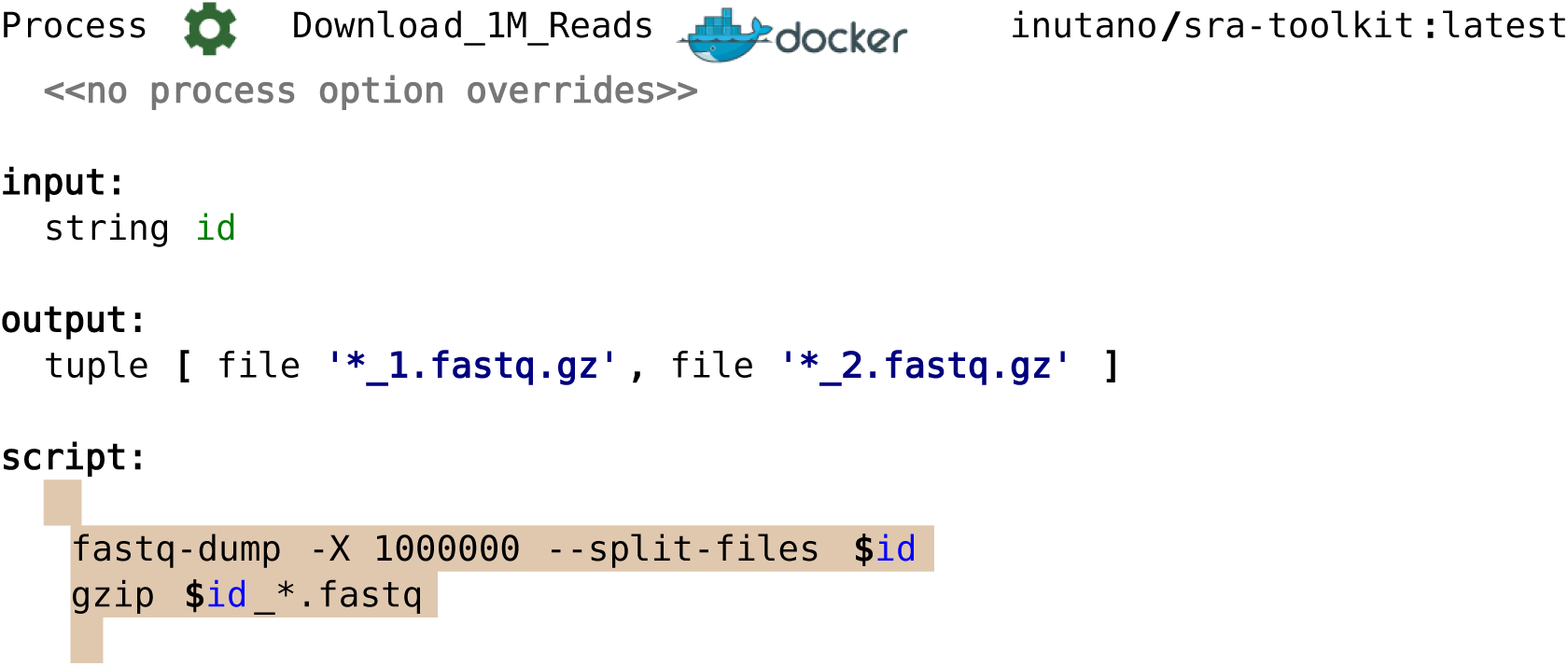
NextflowWorkbench Process. A Process defines inputs, outputs, an optional docker container, and a script. In the example shown, the process accepts an input called ‘id’ of type string. The string is used to query the Short Read Archive with the sra-toolkit and retrieve paired FASTQ files. Inputs must be available before the script can execute. Similarly to Nextflow, a process only executes successfully if the outputs it declared have been produced by the script execution. Notice how the Process incorporates graphical elements and colors to clearly mark different roles of the language elements. When a docker container is specified, as shown in this figure, the commands shown in the script will be executed inside the container. This semantic is implemented by the Nextflow execution runtime. NextflowWorkbench provides autocompletion for input arguments inside the script. This mechanism reduces the risk of typos in variable names and provides instant refactoring of variable names across the script when the workflow programmer renames an input variable.

In contrast to Nextflow, Processes in the NextflowWorkbench are created independently from a Workflow script (i.e., outside the script). As standalone language constructs (implemented as MPS root nodes), Processes can be developed and packaged to be shared with others (for instance as solutions provided in MPS plugins).

As usual when developing a language with LWT, most parts of a Process can be extended by a user of NextflowWorkbench using language composition. The next section describes an application of language composition where we extended the Script part of a Process with the ability to automatically install data resources needed by the script.

### Processes with Variable Data Resources

Docker containers are useful to isolate the process from the machine where the Process executed, but they have limitations. As we developed bioinformatics workflows with the NextflowWorkbench, we found that docker is only a partial solution when a process requires variable data resources.

As an example, consider the process shown in Figure 3. This Process estimates counts for RNA-seq reads against a transcriptome. Different species have different transcriptomes, and different reference transcriptomes or build versions exist for the same species. With the mechanisms provided by docker, one would create different images for different combinations of species and reference build number. This is not really practical because there are a large number of these combinations and only a few may be of practical interest. If a core facility wanted to provide workflows to support multiple species, core personnel would have to anticipate the needs of its user base and configure a large number of possible images to support any species that the core users could need.

**Figure 3.**
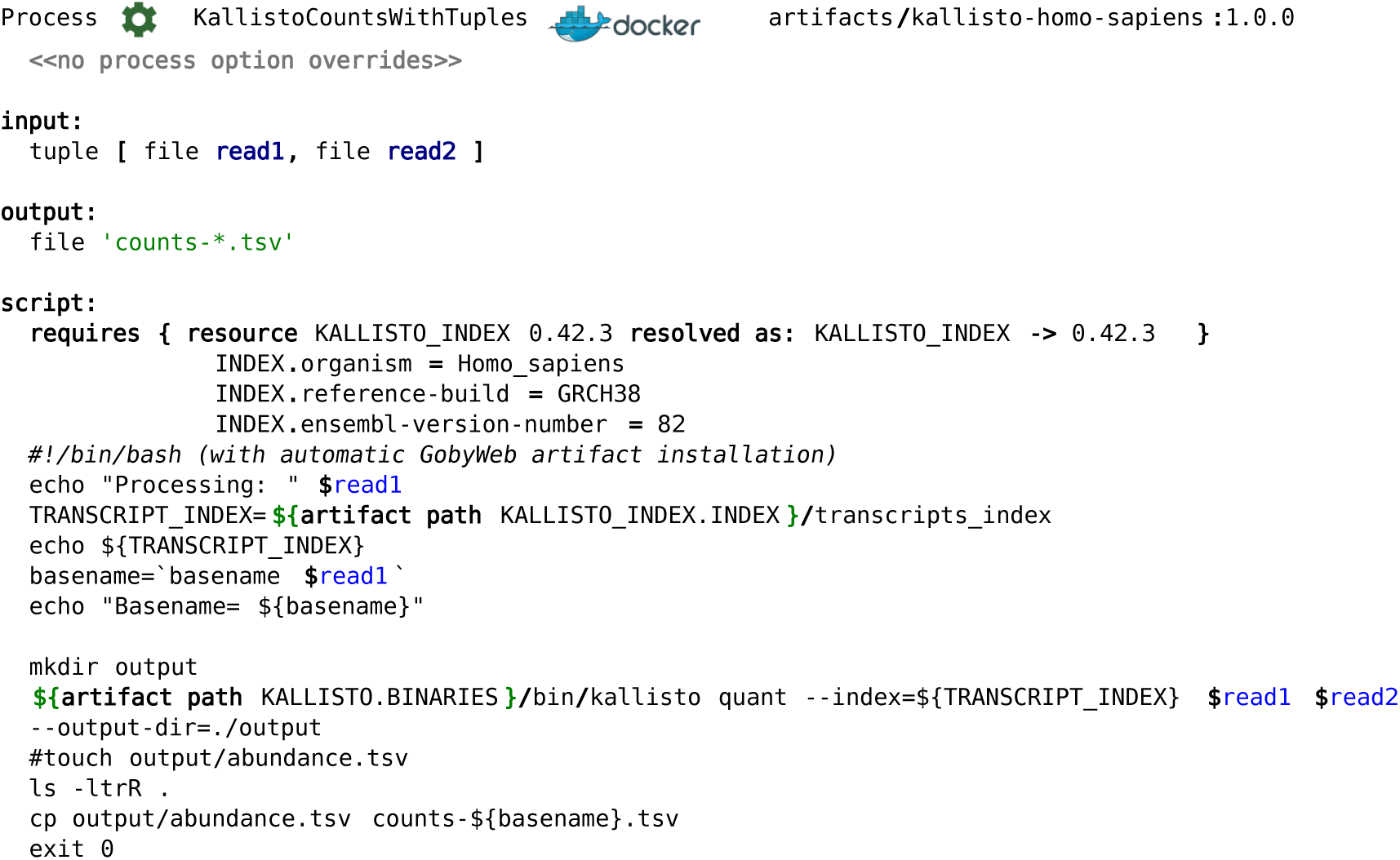
Process with GobyWeb Artifact Installation. This process uses a special type of script which declares dependencies on GobyWeb resources. GobyWeb resources can automatically install variable data resources, such as a specific transcriptome index identifier by species, reference build and Ensembl version number (as shown). In this example, the script requests installation of the `KALLISTO_INDEX` resource version 0.42.3. This resource was configured to retrieve the human transcriptome corresponding to `GRCH38`, in Ensembl version `82`. Notice that rather than writing the complicated steps to download and index this transcriptome, the workflow developer can express the data dependency declaratively.

Rather than creating static images with all the data packaged in a container and for all possible choices of interest, it can be more efficient to provide a mechanism to declare what specific resource is needed, and use software already packaged inside the docker image to assemble data resources on demand. In this case, assembling the transcriptome resource consists in downloading the appropriate transcriptome reference sequence from Ensembl and indexing this reference with the Kallisto program. We have developed mechanisms to support this on-demand strategy, and make it easy to construct specific data resources. The use of these mechanisms is shown in Figure 3, where a simple `requires` block declares a dependency on the `KALLISTO_INDEX` data resource. The same approach also supports choosing and installing specific versions of software resources.

Resource installation scripts are built with the method previously developed for GobyWeb Dorff et al. [2013]. Examples of resources configurations are distributed on GitHub at https://github.com/CampagneLaboratory/gobyweb2-plugins/tree/master/plugins/resources (Campagne et al. [2016]).

### Interactive Docker Features

Developing scripts that run inside a docker container can be challenging because the programmer has to know and remember what programs and data are available inside the container and their precise location. The traditional way to build this understanding is to use interactive console sessions manually started inside a container. The shell can then be used to inspect the files and programs available in the running container and the user has to copy and paste these locations in the script. In NextflowWorkbench, we developed an auto-completion feature that shows container directories interactively when writing the Process script. With this method, a developer can associate a docker image to a Process, start an interactive container, and use auto-completion in the script to locate files or directories of interest (see Figure 4 for an illustration) without even leaving the workbench. Writing correct paths becomes seamless and no longer requires switching between Process editor and console. Similar capabilities are available to auto-complete commands in PATH as well as finding the exact location of data or programs installed as GobyWeb resources (an example is shown in Figure 3 where `$artifact paths` can be assembled using auto-completion).

**Figure 4.**
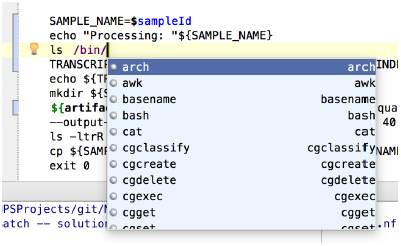
Interactive Path Auto-Completion. This figure illustrates interactive auto-completion of paths inside a docker container. After starting an interactive docker container, Process developers can auto-complete paths inside the docker container in the editor. Similar auto-completion functionality is offered for the GobyWeb data and program resources. These interactive features are implemented with standard features of the MPS language workbench.

### Workflow

A NextflowWorkbench Workflow consists of a set of inputs, references to processes and a list of optional report clauses. Assume that a user wishes to program a workflow to automate the analysis shown in Figure 5. Figure 6 illustrates how such a workflow can be expressed with the NextflowWorkbench language. In contrast to Nextflow, which define Processes inside a workflow script, a NextflowWorkbench Workflow accesses to processes by reference. Process references make it possible to name the Process’ inputs and outputs in the context of the workflow. Such names are used to establish connections between process invocations. For instance, in Figure 6, the reference to `KallistoCountsWithTuples` associates the name `B` to the input of the Process, and associates the name `result` to its output. When the name `result` is defined in this way, the user becomes able to bind the name `result` in the input role of another process reference.

**Figure 5.**
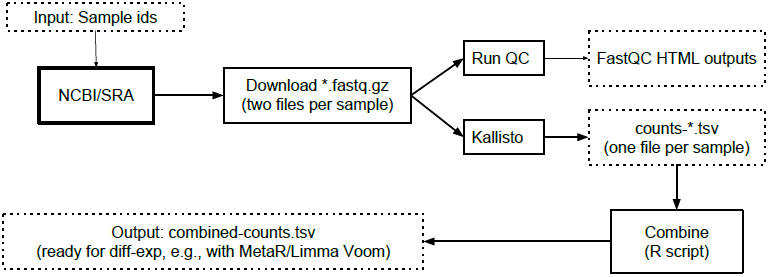
Diagram of Workflow. This diagram is a schematic representation of the analysis workflow shown in Figure 6.

**Figure 6.**
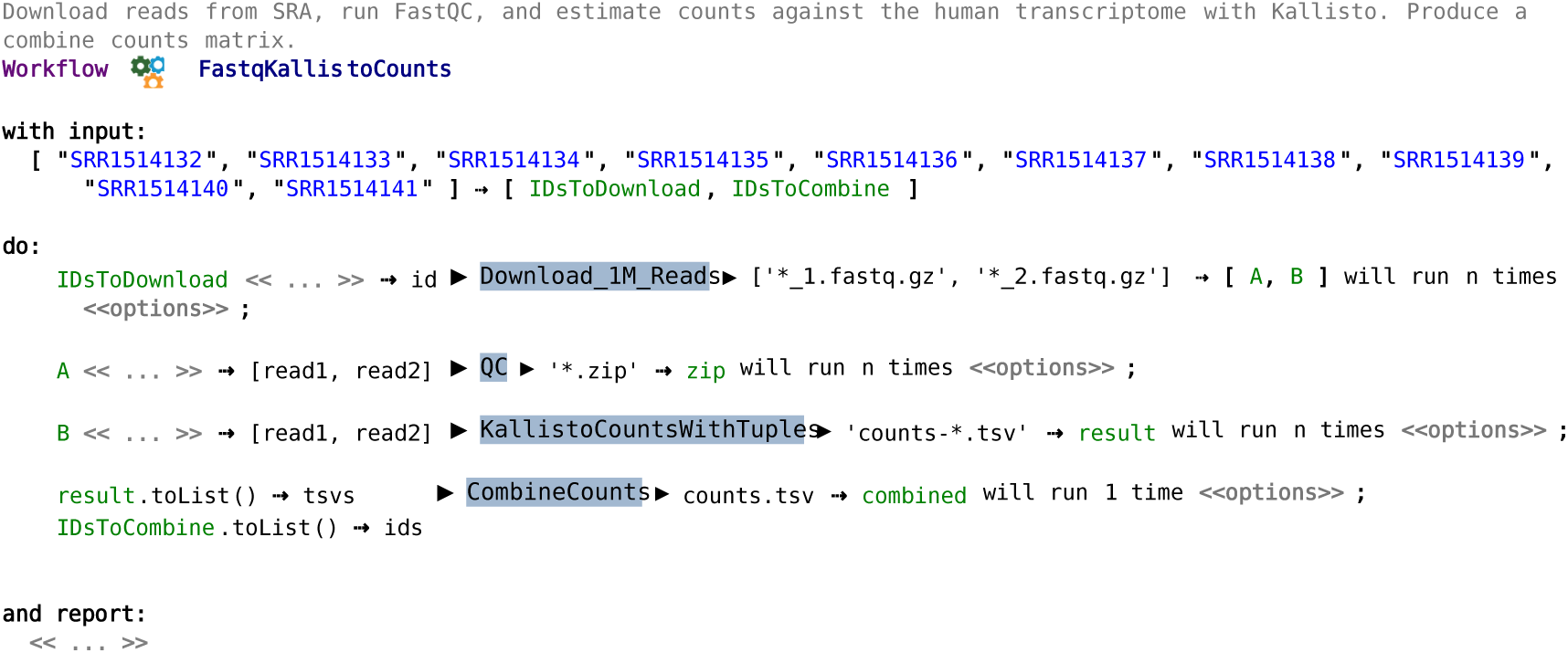
Workflow Example. In this example, a set of SRA identifiers is defined in the input section of the workflow. The list is then duplicated and fed to two processes: `Download_1M_Reads` and `CombineCounts`. The `Download_1M_Reads` will run once for each identifier. When this process terminates, the output, which consists of the tuple of files [’*_1.fastq.gz’, ’*_2.fastq.gz’], is duplicated to be fed to the `QC` and `KallistoCountsWithTuple` processes. The counts obtained with Kallisto are fed to `CombineCounts` along with the list of ids to produce a combined matrix of counts for all the samples analyzed.

In order to prevent cyclic dependencies, output names cannot be set on an input of the same process, and can be set on one input at most. When an output needs to be consumed by several downstream processes (e.g., to implement the fork after Download in Figure 5), the language offers an intention to duplicate a name (an intention is a context dependent menu, see Simi and Campagne [2014]). In Figure 6, the symbols → `[A, B]` indicates that the output of the `Download_1M_Reads` process is duplicated and available through the names `A` and `B`. Note that the editor supports adding additional names between the brackets to replicate the name as many times as needed.

### Utility to Non-Programmers

An important question is whether the user interface provided by Nextflow-Workbench provides sufficient assistance to help non programmers with a biology or clinical background develop workflows.

To address this question we developed training material and started teaching how to develop the workflow shown in Figure 6. In these training sessions, biologists and clinicians develop the `Download_1M_Reads` and `QC` processes and reuse the `KallistoCountsWithTuple` and `CombineCounts` processes from a library. Trainees are emailed instructions (http://campagnelab.org/software/nextflow-workbench/instructions-for-workflow-tutorial/) to install the software on their laptop prior to the training session.

The main challenges we have encountered in these training sessions are related to the installation of the software on the trainees machines. About 30-45 minutes of the training sessions are spent verifying installations and performing some installation steps that the trainees have missed. In few cases, instructors are unable to complete an installation because the trainee computers did not meet minimal specifications (e.g., outdated operating system version, minimum required is Mac OS 10.8.3, or memory requirements not met, minimum needed is 8GB to run Kallisto inside docker on a Mac laptop). Reducing the number of installation steps and provisioning a cluster for remote execution of workflows would go a long way to make this training more accessible and we are actively developing solutions to this end.

We found that we could teach trainees whose laptop met requirements how to assemble the workflow shown in Figure 6 in less than two hours (including time to troubleshoot installations).

### Advanced Docker Features

The NextflowWorkbench can be used as an interactive development environment for developing docker images. We have developed a composable Dockerfile language. Figure 7 illustrates how docker build files can be written in the workbench to assemble a docker image. In this figure, the first two instructions (`FROM` and `MAINTAINER`) will be familiar to docker programmers who have written or read Dockerfiles. The last instruction, however, is not part of the standard Dockerfile language. This instruction was added with language composition to make it easier to install GobyWeb software or data resources inside the image.

**Figure 7.**
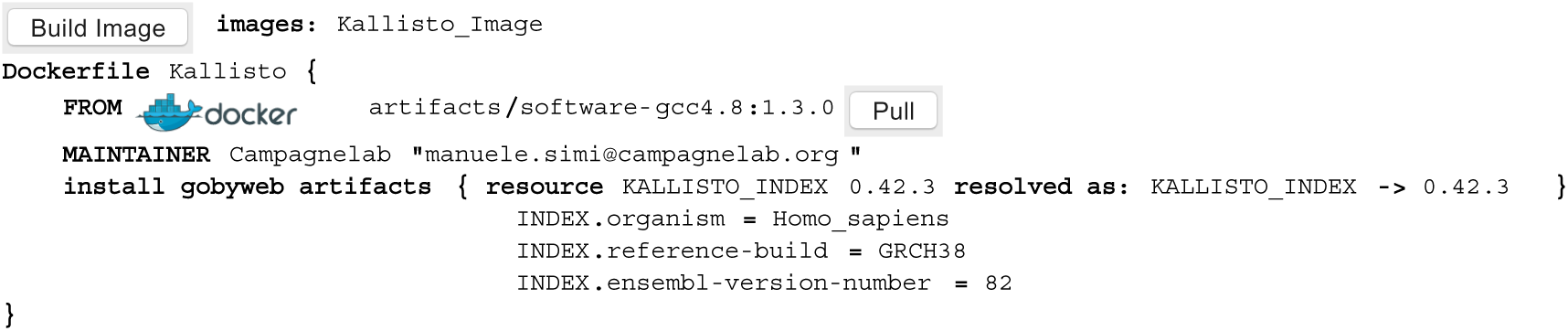
Support for Building Docker Images. NextflowWorkbench offers a composable language to help workflow programmers construct Docker images. In this example, we show a DockerFile root node with a special instruction type called `install gobyweb artifacts`. This special instruction generates `RUN` commands that install GobyWeb resources in the docker image. Pressing the `Build Image` button assembles the image. Built images can be used in Processes. Such instructions are useful to create frozen docker images that contain a specific data resource, for instance for clinical analysis workflows which must be frozen as required by regulations.

**Figure 8.**
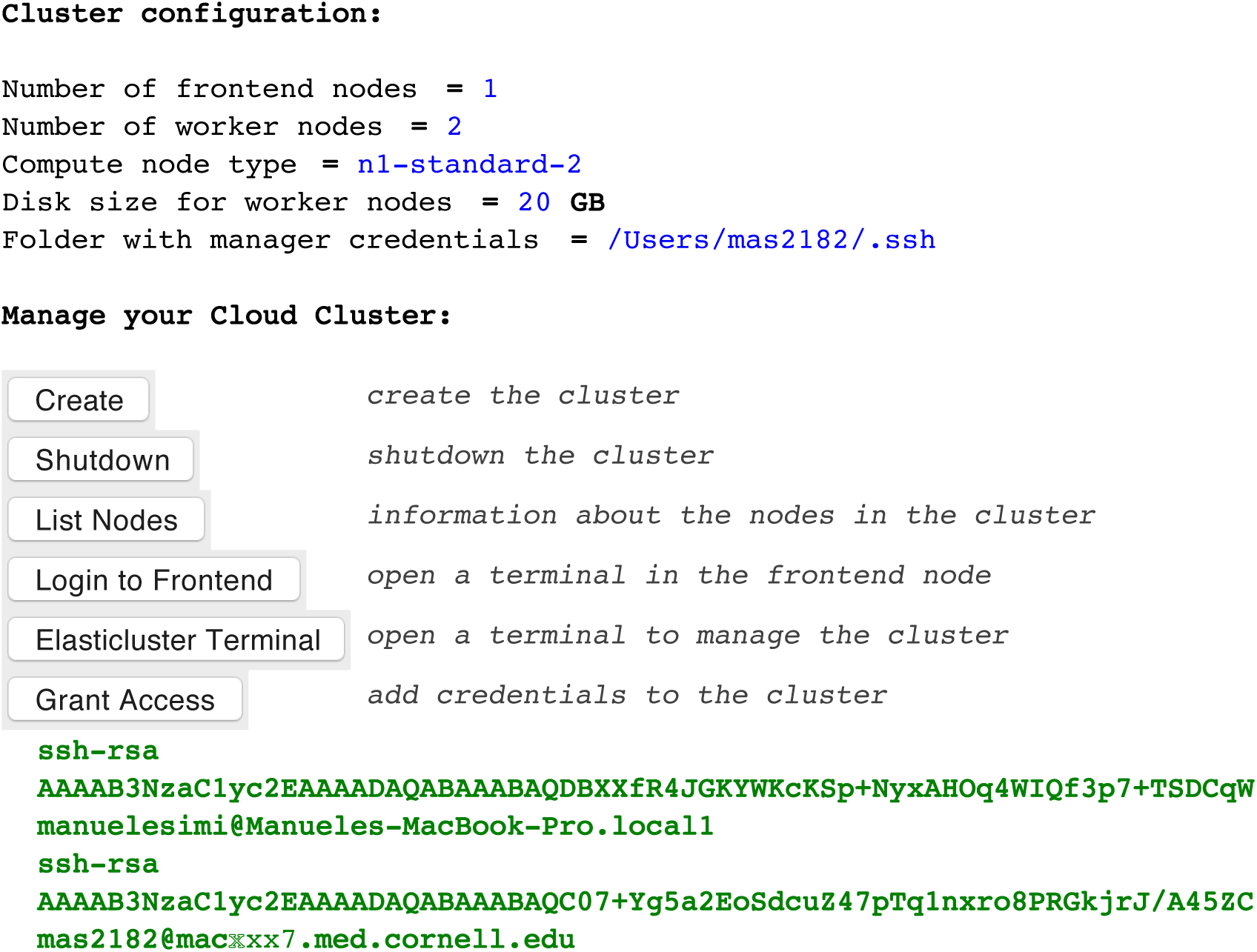
User Interface to Provision a Cluster in the Cloud. This user interface is implemented as an MPS language distributed with NextflowWorkbench. It makes it possible for an instructor to provision a cluster for a training session, or for a laboratory to provision a cluster for a large computation. The language exposes a few parameters of the cluster, and provides actions to create and destroy a cluster, obtain information about the nodes, or login to the cluster front-end. The “Grant Access” button can be used to configure the sets of users who may submit jobs to the provisioned cluster.

### Support for Workflow Execution in the Cloud

Motivated by the difficulty to install docker and run workflows on the trainees laptops, we have experimented with workflow execution in the cloud. A cluster
in the cloud is useful for training because clusters can be created specifically for training sessions and their configurations tuned to a specific type of training. We extended NextflowWorkbench with features to provision a cluster in the Google Cloud. Figure 8 illustrates the user interface available to an instructor, or to members of a laboratory who need the provision a cluster for computation.

## DISCUSSION

Tools to support the development and execution of workflows have been popular among scientists who need to automate repetitive execution of programs to process different inputs. Galaxy and GenePattern are examples of such tools, which we call graphical workflow systems, designed to help biologists who are not programmers take advantage of bioinformatics programs and automate analyses. The tools offer a graphical metaphor for a workflow where boxes represent tools and lines connect the tools where data feeds from one tool to the other. Graphical workflow systems in wide use today were introduced about 10 years ago Giardine et al. [2005], Reich et al. [2006], Hull et al. [2006]. While popularity for these tools grew among computational beginners, experts have yet to widely adopt these systems. While it is unclear what set of reasons explain this lack of interest across the community, understanding these reasons could help design improved systems that both beginners and experts would want to use.

Recent work by others, including Di Tommaso et al. [2014], Cingolani et al. [2015], have developed languages to express workflows with scripting or programming languages in an effort to make these tools more useful to computational experts, including bioinformaticians, or biologists with a programming background. In contrast to graphical workflow systems, scripting workflow systems can be installed very quickly and provide strong support for high-performance computation (including support for implicit or explicit parallelization). The focus of these systems is on helping expert bioinformaticians build high performance parallel data analysis workflows. In addition to the workflow language, these systems offer a runtime system to execute the workflow using a range of high-performance grid schedulers (such as Sun Grid Engine, SLURM or Apache Ignite). Because these systems require expressing workflows as source code, learning how to use them requires becoming familiar with the syntax of a new computational language.

Learning syntax is a difficult part of learning a programming language. For instance, Denny et al. [2011] studied the frequency of syntax errors made by 330 students taking an introductory Java course at the University of Auckland. They found that the top students made syntax errors in 50% of their coding attempts, and that the proportion of students who encountered syntax issues (at least four consecutive program submissions with syntax errors) was 2/3 across most student levels. This clearly illustrates that syntax is often a substantial barrier for novices who learn a new language. NextflowWorkbench offers several interactive features that help novices and guide them while they learn the nextflow syntax. For instance, auto-completion is a powerful feature that can be taught to students quickly and that provides help in every context of a NW workflow. Because the NW language is developed in MPS, rather than in text, it is also much harder to introduce syntax errors by accident. Keywords of the language are treated as constants, which cannot be accidentally modified. Errors are highlighted in red and provide immediate feedback to the workflow programmer. These features help guide new users while they learn the language.

Kronos is a recent workflow system presented in a pre-print Taghiyar et al. [2016]. Kronos supports writing workflows as structured text files that get compiled into python scripts for execution on a variety of computational platforms (local execution, cluster and cloud). The structured file format makes it possible to define tasks (analogous to Nextflow processes) and subsequently connect these tasks using input output I/O connections. The Nextflow language supported by NextflowWorkbench seems more flexible than simple I/O connections because it is possible to transform data produced by processes with a sequence of functions (e.g., see the use of the `toList()` functions in Figure 6) before the data is provided to a process. A large number of pre-defined functions are supported by Nextflow as well as user-defined closures that can be used to process data in custom ways. This capability is not clearly apparent in the version of the Kronos preprint available as of this writing. Kronos also aims to provide a system that both beginners and experts can use, but does not provide a user interface and integrated development environment. Despite its simplicity, the Kronos language represents a new text-based language whose syntax must be learned by beginners who will find out about errors when running the compiler. In contrast, NextflowWorkbench provides an interactive graphical user interface and advanced features, such as real time error detection, on top of an expressive workflow language: Nextflow.

In a broader context, languages to express workflows are essential elements of infrastructures where data analyses are moved to the data. Such infrastructures are becoming necessary in situations where volumes of data are very large (i.e., a tera-byte and more). When several groups need to make computations on the same set of data, transferring the data to the analysis code becomes a bottleneck. In these cases, it is more efficient to move the code to the data than to do the opposite. In the USA, the National Cancer Institute at the National Institutes of Health has started a series of pilots to evaluate this type of infrastructure. The Broad institute, who leads one of these pilot infrastructure projects, has developed the Workflow Description Language and supports executing workflows expressed in WDL on the Broad infrastructure. Another, similar, but incompatible file format to express workflows is the Common Workflow Language (CWL), developed by the Seven Bridges Cancer Genomics Cloud in another NIH/NCI cloud pilot. CWL aims to become a widely used standard to express workflows. To this end, the project organizers are trying to engage a large community of people in the design of CWL. NextflowWorkbench differs from these efforts in several important ways. First, the focus is on user experience to enable beginners to develop and use workflows and to make experts more productive. Neither WDL or CWL address this need. Second, because we used LWT to develop NextflowWorkbench, others can develop extensions of the languages that become immediately integrated with NextflowWorkbench (through language composition and micro-language design, as illustrated in Campagne and Simi [2015]). In our experience, in most cases, there is no need for coordination with our group to develop simple extensions and share them with others. This differs strikingly from a standard development effort, which requires numerous formal communications and coordination before any change can be made to the specification of the “standard”.

Execution of workflows in the cloud is a useful feature for training sessions or when the volume of analyses grows beyond the capability of a laboratory compute resources. While the feature is useful, provisioning a cluster in the cloud can be far from trivial. For instance, the Google Genomics project provides instructions for creating a cluster with elasticluster, which requires an instructor to install tools and dependencies before configuring several text files (http://googlegenomics.readthedocs.org/en/latest/use_cases/setup_gridengme_cluster_on_compute_engine/). Completing these steps and configuring elasticluster may require weeks of effort. In contrast, NextflowWorkbench provides a seamless interface to provision a cluster in a commercial cloud. This interface is constructed with a language that adds this functionality to the MPS user interface, and with docker containers that provide the necessary software dependencies.

## CONCLUSION

In this study, we presented the design and implementation of a workflow system meant for a broad spectrum of potential users, ranging from computational beginners to expert bioinformaticians. We applied language workbench technology to develop such a system, using Nextflow as underlying workflow execution system. We found that we could successfully teach this new workflow system to non-programmers who are able to develop and reuse the simple workflow presented in this manuscript in a short training session of two hours. The NextflowWorkbench platform supports workflow development from laptop computers to clusters provisioned on a commercial cloud.

## ACKNOWLEDGMENTS

The authors thank Paolo Di Tommaso for assistance integrating changes needed for the NextflowWorkbench into Nextflow distributions. We thank the members of Cedric Notredame’s Laboratory (http://www.crg.eu/cedric_notredame) for developing and maintaining Nextflow as an open-source project. This investigation was supported by the National Institutes of Health NIAID award 5R01AI107762-02 to Fabien Campagne and by grant UL1 RR024996 (National Institutes of Health (NIH)/National Center for Research Resources) of the Clinical and Translation Science Center at Weill Cornell Medical College. We thank the training session participants who have provided feedback on earlier versions of NextflowWorkbench and help make this new workflow system relevant to computational beginners.

## REFERENCES

V. M. Benson and F. Campagne. Language workbench user interfaces for data analysis. PeerJ, 3:e800, 2015.

F. Campagne. The MPS Language Workbench, volume I. Fabien Campagne, 2014.

F. Campagne. The MPS Language Workbench, volume II. Fabien Campagne, 2015.

F. Campagne. and M. Simi. MetaR Documentation Booklet. Fabien Campagne, 2015.

F. Campagne, W. E. Digan, and M. Simi. MetaR: simple, high-level languages for data analysis with the Recosystem. bioRxiv, page 030254, 2015. doi: 10.1101/030254. URL http://biorxiv.org/lookup/doi/10.1101/030254.

F. Campagne, manuelesimi, and nchambwe. gobyweb2-plugins: Gobyweb plugins for nextflowworkbench manuscript, 2016. URL http://dx.doi.org/10.5281/zenodo.48271.

P. Cingolani, R. Sladek, and M. Blanchette. Bigdatascript: a scripting language for data pipelines. Bioinformatics, 31(1):10–16, 2015.

P. Denny, A. Luxton-Reilly, E. Tempero, and J. Hendrickx. Understanding the syntax barrier for novices. In Proceedings ofthe 16th annual joint conference on Innovation and technology in computer science education, pages 208–212. ACM, 2011.

P. Di Tommaso, M. Chatzou, P. P. Baraja, and C. Notredame. A novel tool for highly scalable computational pipelines. 2014. URL http://dx.doi.org/10.6084/m9.figshare.1254958.

P. Di Tommaso, E. Palumbo, M. Chatzou, P. Prieto, M. L. Heuer, and C. Notredame. The impact of docker containers on the performance of genomic pipelines. PeerJ, 3:e1273, 2015.

S. Dmitriev. Language oriented programming: The next programming paradigm, 2004. URL http://www.onboard.jetbrains.com/is1/articles/04/10/lop/.

K. C. Dorff, N. Chambwe, Z. Zeno, M. Simi, R. Shaknovich, and F. Campagne. GobyWeb: Simplified Management and Analysis of Gene Expression and DNA Methylation Sequencing Data. PLoS ONE, 8 (7), 2013. ISSN 19326203. doi: 10.1371/journal.pone.0069666.

B. Giardine, C. Riemer, R. C. Hardison, R. Burhans, L. Elnitski, P. Shah, Y. Zhang, D. Blankenberg, I. Albert, J. Taylor, et al. Galaxy: a platform for interactive large-scale genome analysis. Genome research, 15(10):1451–1455, 2005.

D. Hull, K. Wolstencroft, R. Stevens, C. Goble, M. R. Pocock, P. Li, and T. Oinn. Taverna: a tool for building and running workflows of services. Nucleic acids research, 34(suppl 2):W729–W732, 2006.

S. M. Kurs, Jason P. and F. Campagne. NextflowWorkbench Documentation Booklet. Fabien Campagne, 2015. URL https://play.google.com/store/books/details/Jason_P_Kurs_Nextflow_Workbench_Documentation_Book?id=VQhVCgAAQBAJ.

M. Reich, T. Liefeld, J. Gould, J. Lerner, P. Tamayo, and J. P. Mesirov. Genepattern 2.0. Nature genetics, 38(5):500–501, 2006.

M. Simi.and F. Campagne. Composable languages for bioinformatics: the nyosh experiment. PeerJ, 2014. URL https://peerj.com/articles/241/.

M. J. Taghiyar, J. Rosner, D. Grewal, B. Grande, R. Aniba, J. Grewal, P. C. Buotros, R. D. Morin, A. Bashashati, and S. Shah. Kronos: a workflow assembler for genome analytics and informatics. Technical report, feb 2016. URL http://biorxiv.org/content/early/2016/02/19/040352.abstract.

M. Voelter. Integrating prose as first-class citizens with models and code. In MPM@ MoDELS, pages 17–26. Citeseer, 2013.

M. Voelter, D. Ratiu, B. Schaetz, and B. Kolb. mbeddr: an extensible c-based programming language and ide for embedded systems. In Proceedings ofthe 3rd annual conference on Systems, programming, and applications: software for humanity, pages 121–140. ACM, 2012.

M. Wilde, M. Hategan, J. M. Wozniak, B. Clifford, D. S. Katz, and I. Foster. Swift: A language for distributed parallel scripting. Parallel Computing, 37(9):633–652, 2011.

